# Gene expression profiling of *corpus luteum* reveals the importance of immune system during early pregnancy in domestic sheep

**DOI:** 10.1101/673558

**Authors:** Kisun Pokharel, Jaana Peippo, Melak Weldenegodguad, Mervi Honkatukia, Meng-Hua Li, Juha Kantanen

**Author notes:** Correspondence: MHL,; JK.

## Abstract

The majority of pregnancy loss in ruminants occurs during the preimplantation stage, which is thus the most critical period determining reproductive success. While ovulation rate is the major determinant of litter size in sheep, interactions among the conceptus, *corpus luteum* and endometrium are essential for pregnancy success. Here, we performed a comparative transcriptome study by sequencing total mRNA from corpus luteum (CL) collected during the preimplantation stage of pregnancy in Finnsheep, Texel and F1 crosses, and mapping the RNA-Seq reads to the latest Rambouillet reference genome. A total of 21,287 genes were expressed in our dataset. Highly expressed autosomal genes in the CL were associated with biological processes such as progesterone formation (*STAR, CYP11A1*, and *HSD3B1*) and embryo implantation (eg. *TIMP1, TIMP2* and *TCTP*). Among the list of differentially expressed genes, a group of sialic acid-binding immunoglobulin (Ig)-like lectins (Siglecs), solute carriers (*SLC13A5, SLC15A2, SLC44A5*) and chemokines (*CCL5, CXCL13, CXCL9*) were upregulated in Finnsheep, while four multidrug resistance-associated proteins (MRPs) were upregulated in Texel ewes. A total of 17 genes and two non-coding RNAs (ncRNA) were differentially expressed in breed-wise comparisons owing to flushing diet effect. Moreover, we report, for the first time in any species, several genes that are active in the CL during early pregnancy (including *SIGLEC13, SIGLEC14, SIGLEC6, MRP4*, and *CA5A*). Importantly, functional analysis of differentially expressed genes suggested that Finnsheep have a better immune system than Texel and that high prolificacy in Finnsheep might be governed by immune system regulation.

## 1. Introduction

Litter size, a key determinant for the profitability of sheep production systems, is highly dependent on ovulation rate and embryo development in the uterus. Earlier studies have shown that the trait of high prolificacy can result due to the action of either a single gene with a major effect, as in the Chinese Hu, Boorola Merino, Lacaune and small-tailed Han breeds (Mulsant et al., 2001; Souza et al., 2001; Davis et al., 2002, 2006; Chu et al., 2007; Drouilhet et al., 2013), or different sets of genes, as in the Finnsheep and Romanov breeds (Ricordeau et al., 1990; Xu et al., 2018). The Finnsheep or Finnish landrace, one of the most highly prolific breeds, has been exported to more than 40 countries to improve local breeds, although the heritability of ovulation rate is low (Hanrahan and Quirke, 1984). In recent years, a FecG^F^ (V371M) mutation in gene *GDF9* has been identified to be strongly associated with litter size in Finnsheep and breeds such as the Norwegian White Sheep, Cambridge and Belclare breeds, which were developed using Finnsheep (Hanrahan et al., 2004; Våge et al., 2013; Mullen and Hanrahan, 2014; Pokharel et al., 2018).

The success of pregnancy establishment in sheep and other domestic ruminants is determined at the preimplantation stage and involves coordination among pregnancy recognition, implantation and placentation, in which the corpus luteum (CL) and endometrium play vital roles (Geisert et al., 1992; Spencer et al., 2004b, 2007). The preimplantation stage of pregnancy is the most critical period in determining the litter size because of the high embryo mortality during this period. It has been shown that most embryonic deaths occur before day 18 of pregnancy in sheep (Quinlivan et al., 1966; Bolet, 1986; Rickard et al., 2017). However, due to the biological complexity of the process and to technical difficulties, embryo implantation is still not well understood.

The CL is an endocrine structure whose main function is to synthesize and secrete the hormone progesterone. Progesterone production is essential for the establishment of pregnancy. However, if pregnancy is not established, the CL will regress as a result of luteolysis, and a new cycle will begin.

The whole-transcriptome profiling approach enables a deeper understanding of the functions of the CL, which may allow the identification of genes and markers that are differentially expressed, for example, between breeds showing different litter size phenotypes. Although most of the studies associated with early pregnancy have been performed in sheep (Spencer et al., 2004b, 2007; Mamo et al., 2012; Bazer, 2013; Raheem, 2017), only a few studies have applied transcriptomic approaches to the CL. A microarray-based transcriptomic study conducted by Gray et al. (2006) identified a number of genes regulated by progesterone (from the CL) and interferon tau (*IFNT*; from the conceptus) in pregnant vs uterine gland knockout (UGKO) ewes (Gray et al., 2006). In a more comprehensive study conducted by Brooks et al. (2016), transcriptome analysis of uterine epithelial cells during the peri-implantation period of pregnancy identified various regulatory pathways and biological processes in sheep (Brooks et al., 2016). Moore et al. (2016) combined gene expression data with genome-wide association studies (GWASs) to understand the roles of CL and endometrium transcriptomes in dairy cattle fertility (Moore et al., 2016). Another study by identified differentially expressed genes (DEGs) between Day 4 and Day 11 in the CL in cattle (Kfir et al., 2018). Though these studies have certainly enhanced our understanding of the roles of the CL during early pregnancy and in ruminant fertility in general, none of the studies have conducted specific comparisons between breeds with different reproductive potential. Thus, in this study, a comparison of transcriptome profiles between two breeds (high prolific Finnsheep and low-prolific Texel) was conducted to provide insight into the similarities in developmental events in early pregnancy between the breeds. Using F1 crosses we were able to better understand the heritability of the genetic markers. Here, the main goal of this study was to build a global picture of transcriptional complexity in Cl and examine differences in developmental profiles during early pregnancy in sheep breeds showing contrasting fertility phenotypes. Thus, this study has relevance to sheep breeding towards achieving improved reproductive capacity.

## 2. Materials and Methods

### 2.1. Experimental design

All procedures for the experiment and sheep sampling were approved by the Southern Finland Animal Experiment Committee (approval no. ESAVI/5027/04.10.03/2012). The animals were kept at Pusa Farm in Urjala, located in the province of Western Finland, during the experimental period. A total of 31 ewes representing three breed groups (Finnsheep (n=11), Texel (n=11) and F1 crosses (n=9) were included in the main experiment (please note that only 18 of the 31 ewes have been included in this study). Analyses were conducted for two different time points during the establishment of pregnancy: the follicular growth phase (Pokharel et al., 2018) and early pregnancy prior to implantation (current study). After ovary removal, the ewes were mated using two Finnsheep rams, and the pregnant ewes were slaughtered during the preimplantation phase of the pregnancy when the embryos were estimated to be one to three weeks old (Supplementary Table S1). At the slaughterhouse, a set of tissue samples (the pituitary gland, a CL, oviductal and uterine epithelial cells, and preimplantation embryos) were collected and stored in RNAlater reagent (Ambion/Qiagen, Valencia, CA, USA) following the manufacturer’s instructions. Of the collected tissue samples, CL was subjected to current study. One of the CLs was dissected from each ovary. For the present study, and particularly for the RNA-Seq of the CL, six ewes each from the Finnsheep, Texel and F1 cross groups were included. Therefore, out of 31 ewes that were originally included in the main experiment, only 18 have been considered here. The experimental design have been described in more detail in an earlier study (Pokharel et al., 2018).

### 2.2. Library preparation and sequencing

RNA was extracted from the tissues using an RNeasy Plus Mini Kit (Qiagen, Valencia, CA, USA) following the manufacturer’s protocol. The details on RNA extraction have been described previously (Pokharel et al., 2018). RNA quality (RNA concentration and RNA integrity number) was measured using a Bioanalyzer 2100 (Agilent Technologies, Waldbronn, Germany) before sending the samples to the Finnish Microarray and Sequencing Center, Turku, Finland, where library preparation and sequencing were performed. RNA libraries were prepared according to the Illumina TruSeq® Stranded mRNA Sample Preparation Guide (part # 15031047) which included poly-A selection step. Unique Illumina TruSeq indexing adapters were ligated to each sample during an adapter ligation step to enable pooling of multiple samples into one flow cell lane. The quality and concentrations of the libraries were assessed with an Agilent Bioanalyzer 2100 and by Qubit® Fluorometric Quantitation, Life Technologies, respectively. All samples were normalized and pooled for automated cluster preparation at an Illumina cBot station. High-quality libraries of mRNA were sequenced with an Illumina HiSeq 2000 instrument using paired-end (2×100 bp) sequencing strategy.

### 2.3. Data preprocessing and mapping

The raw reads were assessed for errors and the presence of adapters using FastQC v0.11.6 (Simon Andrews). As we noticed the presence of adapters, Trim Galore v0.5.0 (Felix Krueger; Martin, 2011) was used to remove the adapters and low-quality reads and bases. Clean RNA-Seq reads were aligned to the latest sheep reference genome using STAR v2.6.1a (Dobin et al., 2013). The reference genome (oar_rambouillet_v1.0) and annotation (NCBI Ovis aries Annotation Release 103) were used to construct “Star genome” prior to mapping step. In order to facilitate the differential expression analysis, “*--quantMode GeneCounts*” option was included in the STAR mapping command. Alternatively, we performed transcript quantification under the quasi-mapping-based mode in Salmon v1.1.0 (Patro et al., 2017) transcriptome index built using oar_rambouillet_v1.0 transcriptome (https://www.ncbi.nlm.nih.gov/assembly/GCF_002742125.1/).

### 2.4. Differential gene expression analysis

The raw counts quantified as part of STAR mapping were considered for gene expression analysis whereas the Salmon-based transcript estimates (TPM, transcripts per million) were summarized to gene level estimates to prepare a table of genes with their abundance using tximport v1.12.3 (Soneson et al., 2016) Bioconductor package. Prior to gene level summarization, customized “*tx2gene*” data frame was created using the annotation (.*gtf file*) of rambouillet v1.0 transcriptome. We used DESeq2 (Love et al., 2014) for differential gene expression analysis. Transcripts with less than 5 read counts were discarded and technical replicates (samples representing same animal) were collapsed/merged before running the DESeq command. Pairwise differential expression analysis was performed between Finnsheep, Texel and F1 crosses. Differentially expressed genes with adjusted p-value of 0.05 (*padj<0.05*) and absolute log 2(fold change) of 1.5 (*abs(log2FoldChange)>1.5*) were regarded as significant in this study. In DESeq2, Benjamini & Hochberg’s method is used for estimating the adjusted p-values. For identifying a subset of genes potentially differentially expressed due to flushing diet effect, we employed a separate design by including diet as a secondary factor. The statistical analyses were performed in R v3.6.0.

### 2.5. Functional analysis of differentially expressed genes

The ClueGO v2.5.5 (Bindea et al., 2009) plugin in Cytoscape v3.7.0 (Shannon et al., 2003) was employed for gene functional analysis. Prior to performing the analyses, we downloaded the latest versions of the Kyoto Encyclopedia of Genes and Genomes (KEGG) pathways and Gene Ontology (GO) terms. The enrichment analysis was based on a one-sided hypergeometric test with the Benjamini-Hochberg correction method. We used a custom reference set that included a list of all the expressed genes in our dataset. We also modified the default GO and pathway selection criteria in such a way that a minimum of three genes and four percent of genes from a given GO or KEGG pathway should be present in the query list. Furthermore, GO terms with a minimum level of three and a maximum level of eight were retained. Finally, GO terms and/or KEGG pathways sharing similar genes were grouped together using kappa statistics with kappa score threshold of 0.4.

## 3. Results

### 3.1. Phenotypic observations

After removal of the remaining ovary, we counted the number of CLs visually in each animal. With an average of 4.09, Finnsheep had the highest number of CLs, whereas Texel had an average of 1.7 CLs (Table 1, Supplementary Table S1). F1 showed phenotypes closer to those of Finnsheep than those of Texel, having 3.75 CLs on average (Supplementary Table S1); this was unsurprising, as we observed a similar pattern in an earlier study (Pokharel et al., 2018). We did not observe more than 2 CLs in the Texel or fewer than 3 CLs in Finnsheep or F1. Similarly, on average, Finnsheep had the highest number of embryos (n=2.6), followed by F1 crosses (n=1.8) and Texel (n=1.5). Interestingly, the embryo survival rate in Texel was highest, where 1.5 embryos were present from 1.7 CLs (88%) on average. On the other hand, Finnsheep (63%) and F1 cross (48%) had remarkably low embryo survival rate. While these findings are based on fewer animals, the results are in line with earlier studies (Rhind et al., 1980; Silva et al., 2016) where typically higher litter size is associated with higher embryo mortality and vice versa. It would be of great interest to determine if productivity follows the same pattern in F2 (i.e., F1 x F1) crosses, backcrosses and presumably also in a reciprocal cross.

**Table 1.** Average number of CL and embryos in Finnsheep, Texel and their F1 crosses. The full details of the phenotypes are available in supplementary table S1.

| Breed | # CL | # Embryos |
| --- | --- | --- |
| Finnsheep | 4.1 | 2.6 |
| Texel | 1.7 | 1.5 |
| F1 | 3.75 | 1.8 |

### 3.2. RNA-Seq data

From the 42 libraries (21 from each tissue, including three technical replicates), around 1,104 million (M) raw reads were sequenced, of which 1,094 M clean reads were retained after trimming (Table 2). The summary statistics from Trim Galore revealed that up to 3.6% of the reads were trimmed, with reverse-strand reads having a comparatively higher percentage of trimmed bases. However, the percentage of reads that were excluded for being shorter than 18 bp was always less than 1% across all samples. More than 80% of the reads from all samples were aligned to the Rambouillet reference genome and transcriptome using STAR and Salmon, respectively. The raw RNA-Seq data (Fastq files) from this study are available in European Nucleotide Archive (ENA) under project accession code PRJEB32852. The details of sample summary including ENA accession code of individual samples is available in Table 2.

**Table 2.** Sample Summary. Here, FS, TX and F1 represent Finnsheep, Texel and F1 crosses, respectively. Diet is either normal diet (C) or flushing diet (F). Alignment statistics from STAR include only the uniquely aligned reads whereas those from salmon include all reads that mapped to reference transcriptome.

| Sample | Breed | Diet | Raw reads (M) | Clean reads (M) | Uniquely aligned reads (STAR, M) | Aligned reads (salmon, M) | ENA accession code |
| --- | --- | --- | --- | --- | --- | --- | --- |
| 1033 | TX | C | 56.4 | 55.9 | 48.0 | 50.7 | ERR3349023 |
| 107A | TX | F | 55.0 | 54.5 | 46.5 | 49.3 | ERR3349025 |
| 107B | TX | F | 55.7 | 55.2 | 48.3 | 49.8 | ERR3349027 |
| 251 | TX | F | 41.8 | 41.2 | 35.4 | 36.9 | ERR3349029 |
| 302 | TX | C | 54.5 | 54.1 | 47.1 | 48.5 | ERR3349031 |
| 312A | FS | F | 54.9 | 54.4 | 47.0 | 49.3 | ERR3349033 |
| 312B | FS | F | 79.2 | 78.4 | 69.2 | 71.1 | ERR3349035 |
| 3609 | FS | C | 54.5 | 54.1 | 47.0 | 49.0 | ERR3349037 |
| 379 | TX | F | 43.1 | 42.7 | 36.0 | 38.3 | ERR3349039 |
| 4208 | F1 | C | 54.3 | 53.8 | 44.8 | 49.0 | ERR3349041 |
| 4271A | F1 | F | 47.2 | 46.8 | 40.1 | 42.4 | ERR3349043 |
| 4271B | F1 | F | 46.9 | 46.4 | 40.0 | 42.1 | ERR3349045 |
| 4519 | F1 | F | 51.6 | 51.1 | 44.5 | 46.3 | ERR3349047 |
| 4563 | F1 | F | 54.1 | 53.7 | 45.7 | 48.6 | ERR3349049 |
| 4590 | F1 | F | 48.5 | 48.1 | 40.0 | 43.5 | ERR3349051 |
| 4823 | F1 | C | 52.6 | 52.1 | 42.6 | 47.2 | ERR3349053 |
| 48 | FS | C | 53.4 | 53.0 | 44.7 | 47.9 | ERR3349055 |
| 554 | FS | F | 56.0 | 55.5 | 48.1 | 49.7 | ERR3349057 |
| 73 | TX | C | 46.1 | 45.7 | 39.1 | 41.3 | ERR3349059 |
| 897 | FS | F | 53.3 | 52.9 | 46.1 | 47.9 | ERR3349061 |
| 974 | FS | C | 45.1 | 44.7 | 38.6 | 40.4 | ERR3349063 |

### 3.3. Gene expression in the CL

STAR-based alignment revealed 23,327 genes and pseudogenes expressed in the whole data set which makes up approximately 70% of the list (n=33,372) available for the recent Rambouillet reference genome. Further grouping of the expressed genes showed that the most genes (n=21,052) were expressed in Finnsheep followed by Texel (n=20,957) and F1 crosses (n=20,928). The cumulative difference in the number of genes in different samples and breeds might be due to transcriptional noise. As shown in Fig. 1, the highest number of breed-specific genes was found in Finnsheep (n=399), followed by Texel (n=363) and F1 crosses (n=303). In a pairwise comparison, based on overall gene expression, Finnsheep and F1 crosses shared a higher number of genes (n=338) than other pairs, indicating the closer relatedness of F1 crosses to Finnsheep (Fig. 1). As indicated by the principal component analysis (PCA), we did not observe any breed-specific clusters in either of the tissues and was also the case in our earlier ovarian transcriptome study (Pokharel et al., 2018). Alternatively, salmon-based quantification revealed 27,072 genes expressed in the CL of which 18,463 had TPM greater than 0.1 (Supplementary Table S2).

**Figure 1:**
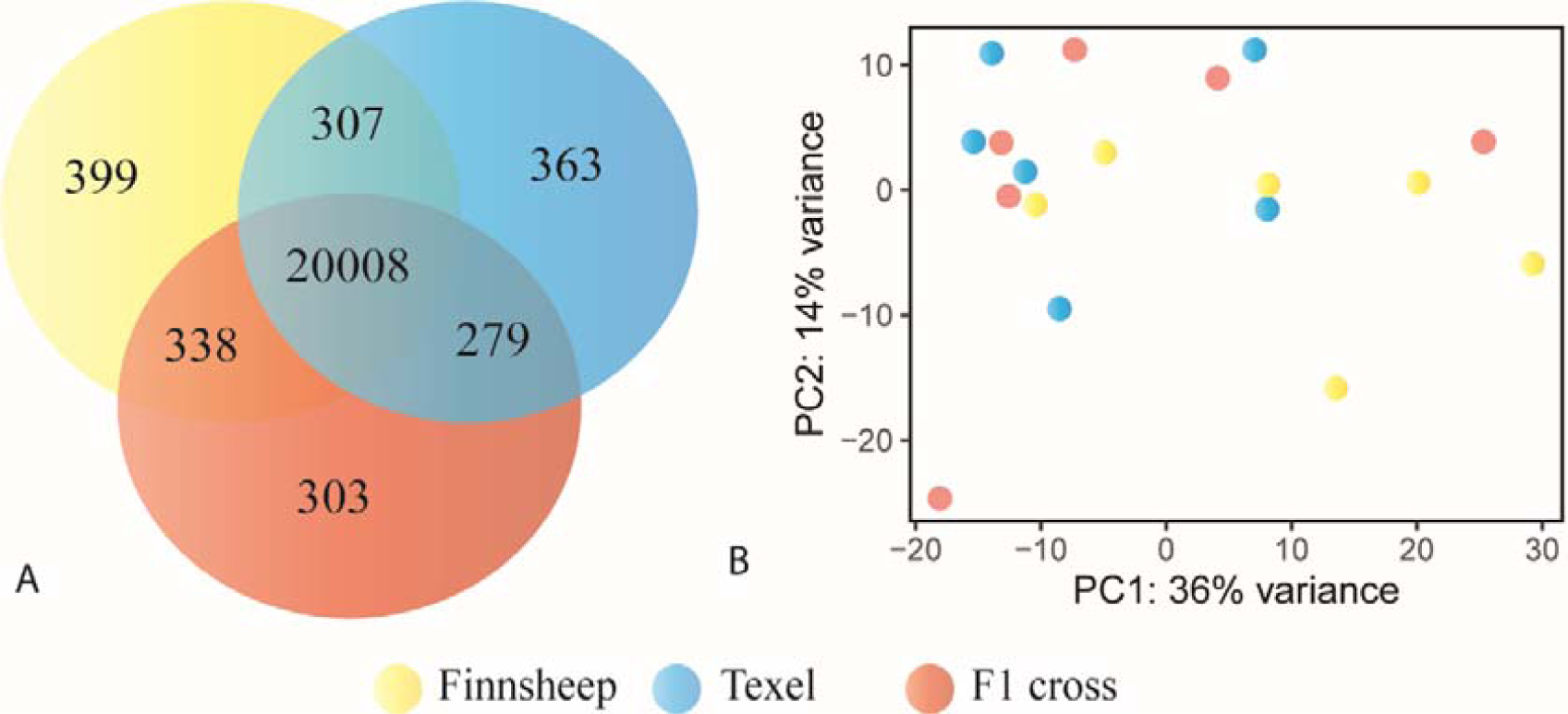
Graphical summary of gene expression in the CL. (A) Shared and unique genes expressed in the CL of Finnsheep, Texel and their F1. (B) PCA plot based on variance stabilized transformation (VST) of gene expression counts derived from DESeq2 in three breeds.

### 3.4. Highly expressed genes

To obtain an overview of the most abundant genes in the tissue, we selected the top 25 genes expressed in the CL (Table 3) derived from Salmon quantification. We noticed that 10 out of the top 25 genes were ribosomal proteins. Moreover, top expressed genes also appeared to play substantial roles during the preimplantation stage. Steroidogenic acute regulatory protein (*STAR*), the second most highly expressed gene, plays an important role in mediating the transfer of cholesterol to sites of steroid production (Stocco, 2000; Christenson and Devoto, 2003). Post ovulation, the expression of the majority of genes associated with progesterone synthesis starts to increase and peaks around the late luteal phase, when the CL has fully matured (Juengel et al., 1995; Devoto et al., 2001; Davis and LaVoie, 2018). *STAR*, together with the cytochrome P450 side chain cleavage (P450cc) complex and 3b-hydroxysteroid dehydrogenase/delta5 delta4-isomerase (*HSD3B1*), are the three most important actors involved in progesterone biosynthesis. *STAR* is involved in transporting free cholesterol to the inner mitochondrial membrane. The P450cc complex, composed of a cholesterol side chain cleavage enzyme (*CYP11A1*), ferredoxin reductase (*FDXR*) and ferredoxin (*FDX1*), converts the newly arrived cholesterol into pregnenolone (King and LaVoie, 2009). Finally, *HSD3B1* helps in converting pregnenolone to progesterone (Hu et al., 2010; Plant et al., 2015; Stouffer and Hennebold, 2015; Davis and LaVoie, 2018). Three of these major genes involved in progesterone synthesis (*STAR, HSD3B1*, and *HSD3B1*) were ranked among the top 25 most highly expressed autosomal genes, while *FDX1* (TPM=972.8) and *FDXR* (TPM=221.5) were also highly expressed.

**Table 3.** List of top 25 most abundant genes expressed in the CL. Gene names and gene descriptions for gene IDs starting with “LOC” were retrieved from NCBI Genbank (https://www.ncbi.nlm.nih.gov/genbank/) and, GeneCards (https://www.genecards.org/), respectively.

| GeneID | Gene Description | Chr | mean TPM |
| --- | --- | --- | --- |
| *TIMP1* | Tissue inhibitor of metalloproteinase 1 | X | 16705.6 |
| *STAR* | Steroidogenic acute regulatory protein | 26 | 14226.6 |
| *OXT* | oxytocin/neurophysin I prepropeptide | 13 | 13113.2 |
| *MGP* | matrix Gla protein | 3 | 12294.0 |
| *LOC114112617* | ncRNA (uncharacterized) | un | 12155.9 |
| *LOC114113966* | translationally controlled tumour protein (*TCTP*) | 3 | 8341.7 |
| *HSD3B1* | hydroxy-delta-5-steroid dehydrogenase, 3 beta- and steroid delta-isomerase 1 | 1 | 8249.3 |
| *LOC101117785* | Apolipoprotein A1 (*APOA1*) | 15 | 8213.2 |
| *LOC101110773* | elongation factor 1-alpha 1 (*EEF1A1*) | 10 | 7564.5 |
| *LOC101103639* | uncharacterized (protein coding) | 3 | 7272.8 |
| *LOC114116158* | thymosin beta-4 pseudogene | 1 | 6689.3 |
| *LOC114118103* | 40S ribosomal protein S29 | 14 | 5599.0 |
| *CYP11A1* | Cytochrome P450 Family 11 Subfamily A Member 1 | 18 | 5096.8 |
| *RPLP1* | ribosomal protein lateral stalk subunit P1 | 7 | 5032.2 |
| *LOC114116632* | 60S ribosomal protein L23a (*RPL23a*) | 10 | 4998.9 |
| *FTH1* | Ferritin heavy chain 1 | 21 | 4498.0 |
| *RPS8* | ribosomal protein S8 | 1 | 4402.1 |
| *LOC105606567* | 60S ribosomal protein L39 | 4 | 4215.9 |
| *MSMB* | microseminoprotein beta | 25 | 4093.6 |
| *RPS24* | ribosomal protein S24 | 13 | 4029.1 |
| *RPS17* | ribosomal protein S17 | 18 | 3950.0 |
| *LOC114108766* | 60S ribosomal protein L17 (*RPL17*) | 4 | 3832.5 |
| *RPS18* | ribosomal protein S18 | 20 | 3657.2 |
| *TIMP2* | Tissue inhibitor of metalloproteinase 2 | 11 | 3587.4 |
| *LOC114114908* | 40S ribosomal protein S27 (*RPS27*) | 5 | 3570.4 |

**Table 4.** List of genes differentially expressed due to the effect of flushing diet in Finnsheep (FS), Texel (TX) and F1 crosses (F1).

| Gene | Gene* | BaseMean | LFC | Padj | Condition |
| --- | --- | --- | --- | --- | --- |
| *COX7C* |  | 101.753584 | 1.93 | 0.03584019 | FS vs TX |
| *LOC114116073* | un_ncRNA | 832.4 | -4.60 | 1.26E-28 | FS vs TX |
| *PAPPA2* |  | 24.4 | -2.27 | 0.00091 | FS vs TX |
| *LOC114118704* | TVP23(p) | 38.8 | -2.51 | 1.70E-06 | FS vs TX |
| *ADAMTS4* |  | 892.8005278 | -1.80 | 0.02409534 | FS vs F1 |
| *SHOX2* |  | 74.00640751 | 1.95 | 0.0200087 | FS vs F1 |
| *HOXD10* |  | 99.29244359 | -2.17 | 0.00279421 | FS vs F1 |
| *SCTR* |  | 87.95488542 | 1.70 | 0.0146455 | FS vs F1 |
| *HOXC11* |  | 24.11123058 | -1.81 | 0.01918401 | FS vs F1 |
| *LOC105611004* | un_ncRNA | 40.93791063 | -2.06 | 0.00279421 | FS vs F1 |
| *LOC114116073* | un_ncRNA | 832.360494 | -4.82 | 5.58E-31 | FS vs F1 |
| *LRP11* |  | 124.9792312 | -1.84 | 0.02732988 | FS vs F1 |
| *LOC114118704* | TVP23(p) | 38.85436938 | -1.92 | 0.00279421 | FS vs F1 |
| *BTN1A1* |  | 22.44434414 | 1.86 | 0.02926403 | FS vs F1 |
| *KCNQ1* |  | 856.9910538 | 1.77 | 0.04977007 | FS vs F1 |
| *HNR1PA1* |  | 81.09239312 | 1.93 | 0.0200087 | FS vs F1 |
| *COX7C* |  | 101.753584 | -1.88 | 0.03094773 | TX vs F1 |
| *PCSK5* |  | 469.246216 | 1.69 | 0.01712337 | TX vs F1 |
| *SCTR* |  | 87.9548854 | 1.79 | 0.00816462 | TX vs F1 |
| *HOXC11* |  | 24.1112306 | -1.92 | 0.00816462 | TX vs F1 |
| *PPM1H* |  | 575.325443 | 1.67 | 0.00684851 | TX vs F1 |
| *HOXA9* |  | 74.2026779 | -1.96 | 0.00816462 | TX vs F1 |
| *RDH10* |  | 2603.9004 | 1.56 | 0.00816462 | TX vs F1 |
| *CA5A* |  | 983.421994 | -1.50 | 0.01137482 | TX vs F1 |

Oxytocin (*OXT*) was one of the most highly expressed genes in the CL. In cyclic ewes, *OXT* secreted from the CL and posterior pituitary is widely known to bind with oxytocin receptor (*OXTR*) from the endometrium to concomitantly release prostaglandin F_2α_ (*PGF*) pulses and induce luteolysis (Flint and Sheldrick, 2004; Spencer et al., 2004a, 2004b; Bazer, 2013). However, for noncyclic ewes, *OXT* plays an important role during peri-implantation and throughout pregnancy (Kendrick, 2000). *OXT* signaling is known to be influenced by progesterone, but the mechanism underlying the regulation is not yet clear due to conflicting findings (Grazzini et al., 1998; Gimpl et al., 2002; Fleming et al., 2006; Bishop, 2013). Meanwhile *OXTR* (TPM=0.2) expression was almost negligible compared to OXT expression.

Two members of tissue-derived matrix metalloproteinase (*MMP*) inhibitors, also referred to as tissue inhibitors of metallopeptidases (TIMPs) were among the top 25 most abundant genes of which *TIMP1* was ranked number 1 and *TIMP2*, number 24. In addition to *TIMP1* and *TIMP2*, at least 20 MMPs (including two isoforms of *MMP9* and *MMP24*) two TIMPs, *TIMP3* (TPM=265) and *TIMP4* (TPM=0.5) were expressed in our samples. The high level of TIMPs indicated successful pregnancies as these inhibitors have low level of expression while their target MMPs are elevated during luteolysis (Curry and Osteen, 2001; Kliem et al., 2007; Nissi et al., 2013). TIMPs have important role in regulation of several processes relevant to uterine physiology including angiogenesis (Moses et al., 1990), cell differentiation (Docherty et al., 1985), and embryo development (Nakanishi et al., 1986). Northern blot analyses in several tissues of sheep showed that both *TIMP1* and *TIMP2* had greatest abundance in CL during early pregnancy (Hampton et al., 1995). Moreover, MMPs and TIMPs have important roles in implantation and that the high level expression of *TIMP1* and *TIMP2* indicated active invasion of trophoblast cells during implantation (Librach et al., 1991; Bai et al., 2005). Another study in bovine oviduct showed that *TIMP1* and *TIMP2* were highly expressed during the time of ovulation (Gabler et al., 2001). Hence, these two genes may play significant role during the whole reproduction process – from ovulation to implantation.

Translationally controlled tumor protein (*TCTP*) is a highly conserved, multifunctional protein that plays essential roles in development and other biological processes in different species (Tuynder et al., 2002; Chen et al., 2007; Brioudes et al., 2010; Li et al., 2011; Branco and Masle, 2019). With a maximum level of expression on Day 5 of pregnancy, this protein has been shown to play a significant role in embryo implantation in mice (Li et al., 2011). Consistent with these earlier studies, *TCTP* appeared to have the highest level of expression during the embryo implantation period. Matrix Gla protein (*MGP*) is a vitamin K-dependent extracellular matrix protein whose expression has been shown to be correlated with development and maturation processes (Zhao and Nishimoto, 1996; Zhao and Warburton, 1997) and receptor-mediated adhesion to the extracellular matrix (Loeser and Wallin, 1992). Several studies have reported that *MGP* is highly expressed in the bovine endometrium (Spencer et al., 1999; Mamo et al., 2012; Forde et al., 2013) and we have shown that it’s also the case with CL. The high level of expression of *MGP* in our study is consistent with the results of earlier studies in which this gene was found to be elevated during the preimplantation stage in sheep (Spencer et al., 1999; Gray et al., 2006) and cattle (Mamo et al., 2012). Similarly, (Casey et al., 2005) reported that *MGP* was significantly upregulated in non-regressed compared to regressed bovine CLs. Our data and supporting results from earlier studies on cattle show that *MGP* is highly expressed in CL during the preimplantation stage and plays important roles in superficial implantation and placentation in sheep.

### 3.5. Breed wise gene expression differences in the CL

Out of the 133 significant DEGs in the CL of pure breeds (i.e Finnsheep *vs* Texel), 90 were upregulated in Finnsheep (Supplementary table S4) and the rest were downregulated. In the list of DEGs, we observed a few cases in which more than one gene from the same family was present. Four isoforms of multidrug resistance-associated protein 4-like (*MRP4*) were upregulated in Texel ewes. Earlier reports have suggested a role of *MRP4* in transporting prostaglandins in the endometrium (Lacroix-Pépin et al., 2011), and *MRP4* has been found to be upregulated in the endometrium in infertile cows compared to fertile cows (Moore et al., 2016). Although there are no reports regarding the existence and roles of *MRP4* in the CL, we speculate that the comparatively lower levels of these prostaglandin (PG) transporters in Finnsheep provide a luteoprotective effect. Five sialic acid-binding Ig-like lectins (Siglecs) including three isoforms of *SIGLEC-14* were upregulated in Finnsheep. Siglecs are transmembrane molecules expressed on immune cells and mediate inhibitory signaling (Varki and Angata, 2006). So far, *SIGLEC-13* has been reported only in nonhuman primates; it was deleted during the course of human evolution (Angata et al., 2004). The importance of Siglecs in immune system regulation has been reviewed elsewhere (Pillai et al., 2012). Siglecs constantly evolve through gene duplication events and may vary between species and even within a species (Cao and Crocker, 2011; Pillai et al., 2012; Bornhöfft et al., 2018). Here we have reported the expression of Siglecs in CL which are known to play a role in the immune response during early pregnancy (preimplantation).

Several cytokines including four chemokines (*CCL5, CCL24, CXCL9*, and *CXCL13*), two interleukin receptors (*IL1RN, IL12RB1*) and two interferons (*ISG20, GVINP1*) were upregulated in Finnsheep. Other genes with more than one member included major histocompatibility complexes (MHCs) (*HA1B, MICB, MICA, BOLA-DQB*0101*, etc.), tripartite motif containing proteins (two isoforms of *TRIM5*, and *TRIM10*), cluster of differentiation factors (*CD4, CD69*), CD300 family molecules (*CD300C*, and *CD300H*). A novel protein coding gene (*LOC114116052*) was significantly upregulated in Texel. Nucleotide blast search of this novel gene’s CDS revealed its sequence similarity with *SENP6* in several mammalian species including transcript variants of sheep *SENP6* (*LOC101118793*, 95% query coverage and 98.7% sequence identity). Therefore, we concluded that *LOC114116052* is one of the isoforms of *SENP6*. All four ribosomal proteins (*MRPS33, RPS3A, RPL17*, and *RPL34*) differentially expressed between the pure breeds were upregulated in Finnsheep. Similarly, 16 of the differentially expressed genes were classified as uncharacterized ncRNAs of which nine were upregulated in Finnsheep.

Out of 90 genes that were significantly upregulated in Finnsheep compared to Texel, 62 were recognized by ClueGO. However, 22 of the recognized genes lacked functional annotation. Thus, enrichment analysis for GO terms and KEGG pathways is based on 40 genes. In the end, only 16 genes passed the selection criteria (see materials and methods section) and were associated with 16 representative terms and pathways (Fig3). It turned out that several of the terms and pathways comprised similar genes and grouping based on common genes revealed four different types of terms and/or pathways. The enriched terms and pathways belonged to four main categories based on the way genes were shared. The representative terms and/or pathways associated with upregulated genes were “negative regulation of viral life cycle”, “Th1 and Th2 cell differentiation”, “chemokine receptor binding” and, “dicarboxylic acid transport”. In summary, genes involved in the immune response were upregulated in Finnsheep CL during early pregnancy.

Altogether, 31 genes were differentially expressed between Finnsheep and F1 crossbred ewes, of which 19 genes were upregulated in Finnsheep (Supplementary table S7).

The lowest number (n=27) of DEGs was observed between Texel and F1 crossbred ewes, with two-third genes being upregulated in F1 (Supplementary table S9).

**Figure 2.**
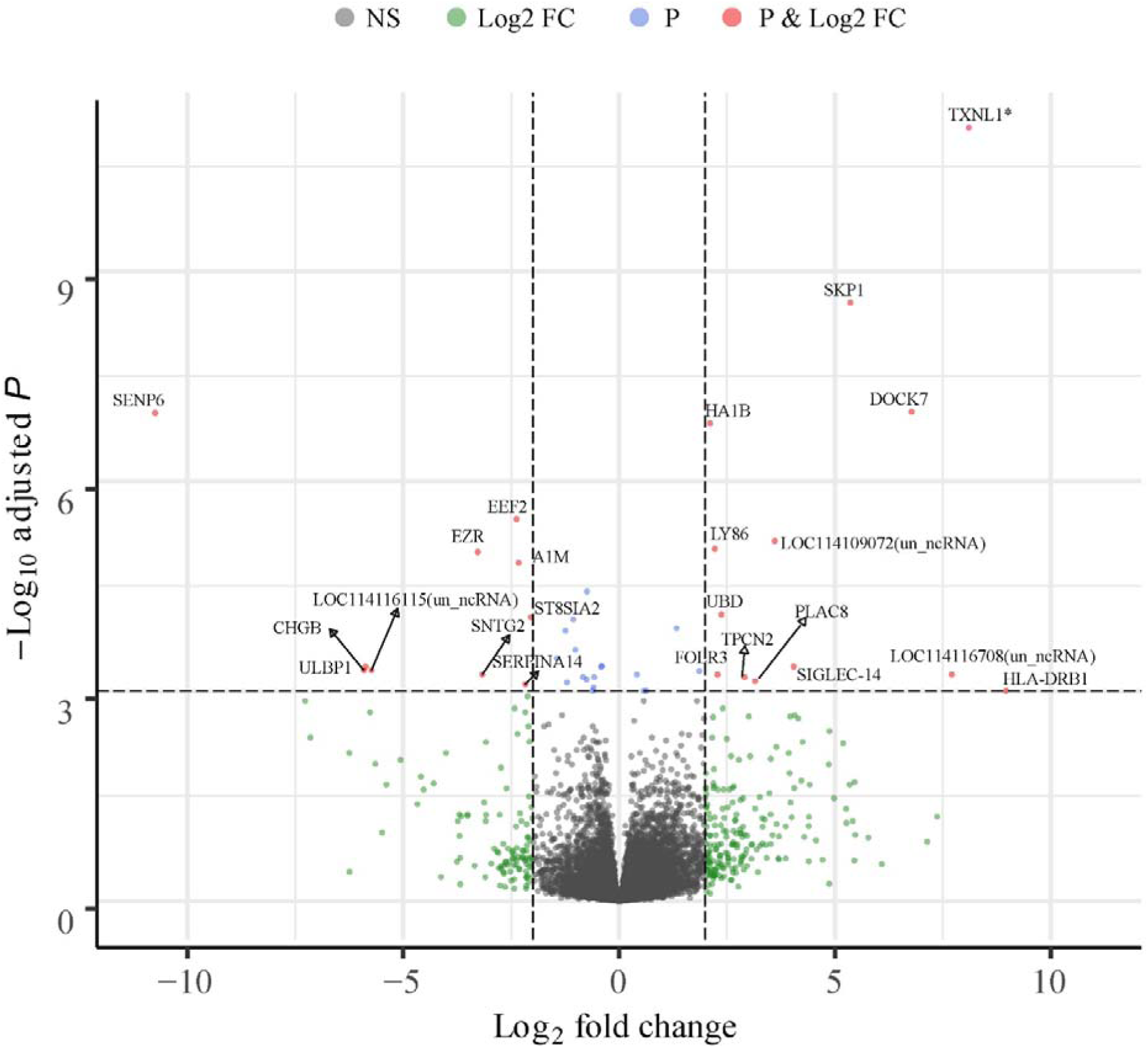
Volcano plot of 23327 genes expressed in the pure breeds. Top 23 of 133 genes significantly differentially expressed between Finnsheep and Texel are denoted by gene names except three unknown ncRNAs. All DE genes with absolute log2 fold change greater than 2 and adjusted P value lower than 0.001 have been highlighted here. For additional details and complete list of 133 DE genes please refer supplementary table. Note that *TXNL1* gene (marked by an asterisk) has an adjusted P-value of 4.84E-16. Volcano plot created using Bioconductor package EnhancedVolcano (Blighe et al., 2019) and some adjustments in the figure were made in Adobe Illustrator (Adobe Inc.). The x-axis represents log2 fold change and the y-axis represents adjusted p-values(-log10). Legends in top: NS = non-significant; Log2 FC = genes having absolute log2 fold change greater than 2; P = genes with adjusted P value less than 0.001; and P & Log2 FC: genes passing the criteria for log2 fold change and adjusted P value.

**Figure 3.**
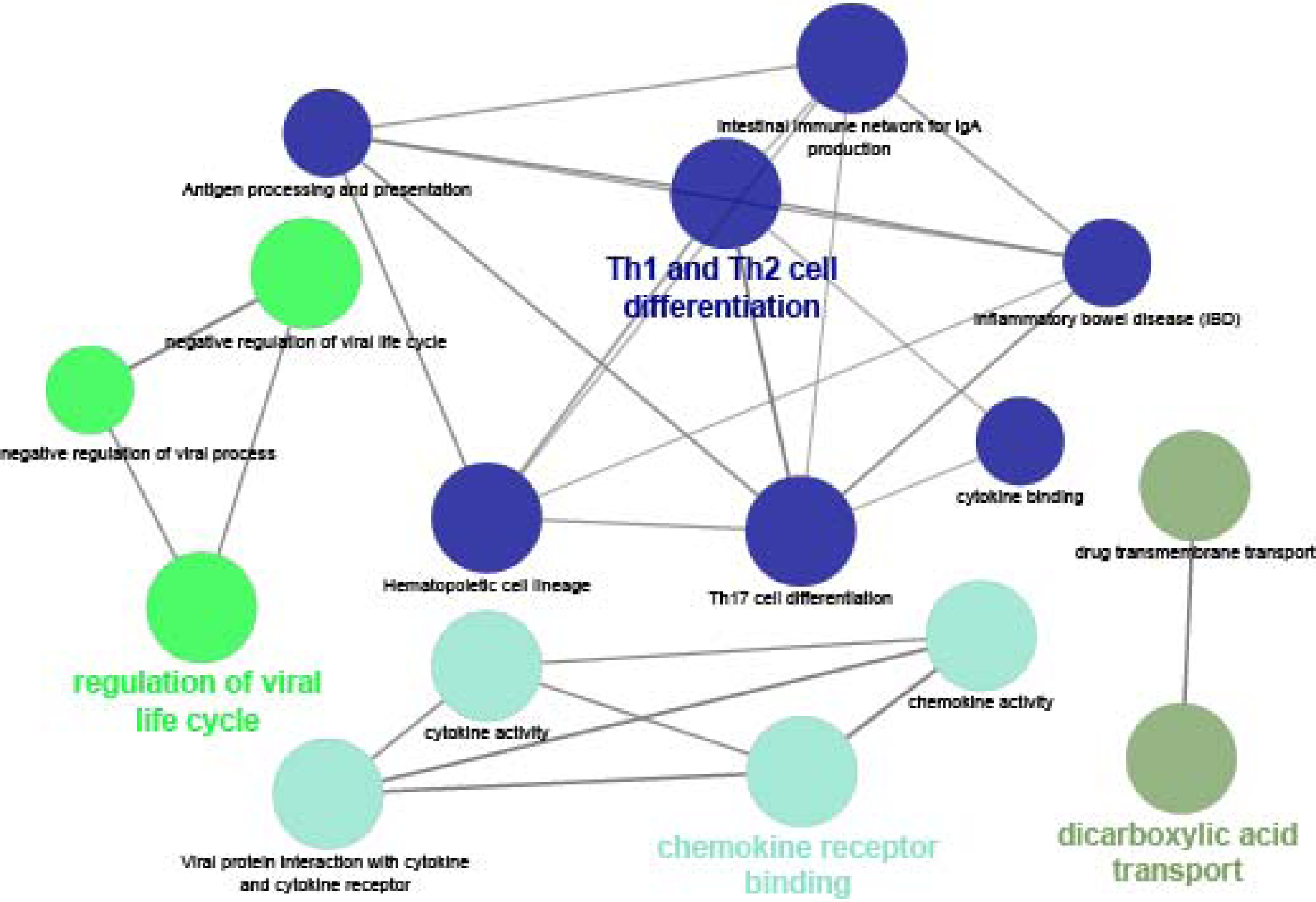
Go terms and KEGG pathways associated with significantly differentially expressed genes between Finnsheep and Texel. The functional grouping option in ClueGO categorized 16 GO terms and pathways into four groups using kappa score where the genes shared between the terms are iteratively compared to form functional groups.

### 3.6. Uniquely differentially expressed genes

We noticed that several genes (see Fig. 4) were differentially expressed in more than one comparison, increasing our confidence in the identification of these DEGs. Few DEGs were exclusively up- or downregulated in particular breed compared to other two breeds. *ANOS1, CLVS2, FOLR3, EEF1A1* and *LOC114115287* (uncharacterized ncRNA) were always upregulated in Finnsheep whereas seven genes were exclusively downregulated. *MEPE* was found to be downregulated in Finnsheep also during the ovulation (Pokharel et al., 2018). Seven genes (*DNER, LOC106990106* (*EZR*), *LOC114109364* (*H3.3*), *LOC114115573* (*COX7C*), *LOC114115623* (*TOMM20*) and *LOC114115666* (*EEF2*)) were exclusively upregulated and 13 genes including three ncRNAs were exclusively downregulated in Texel (Fig. 4).

**Figure 4.**
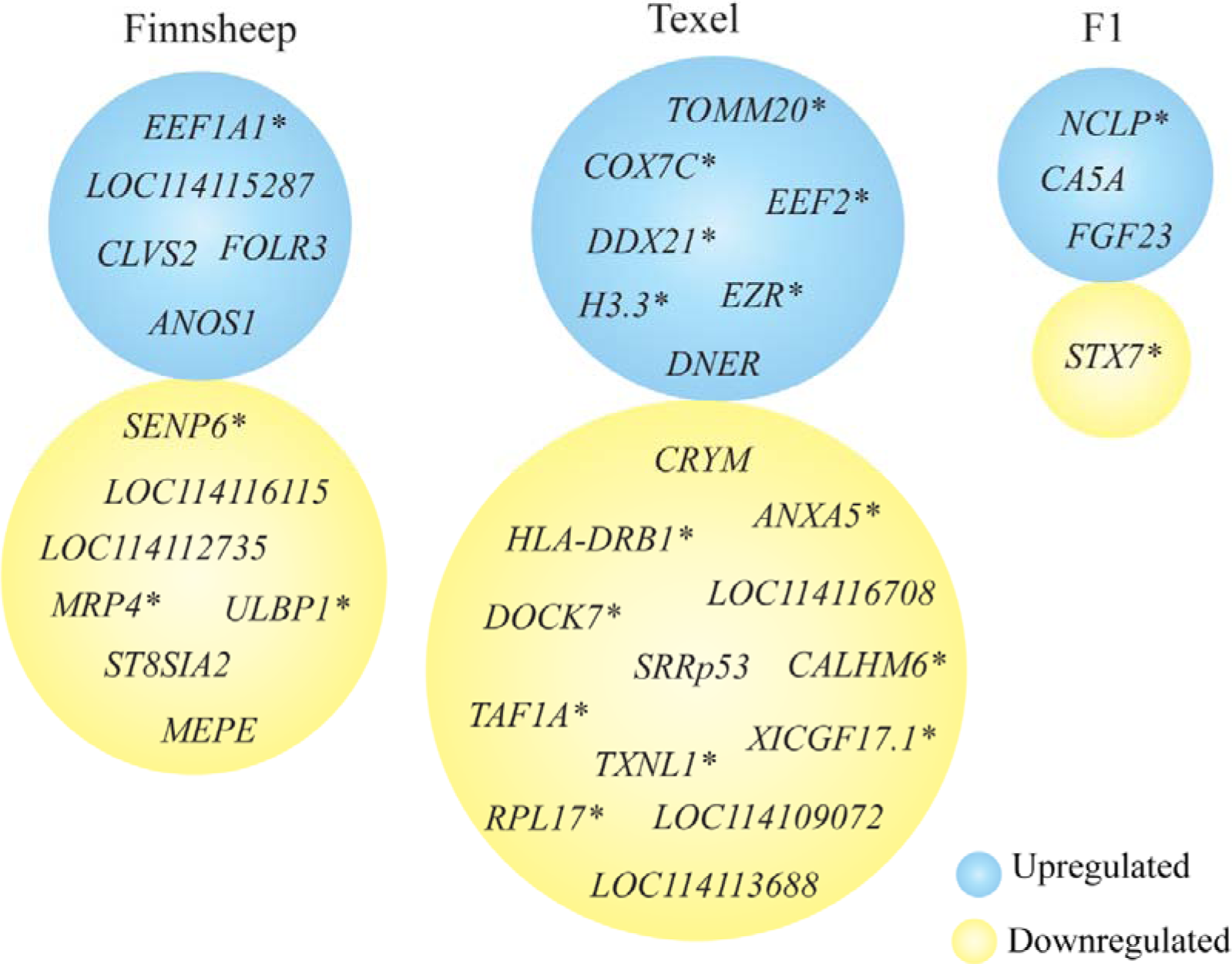
List of uniquely differentially expressed genes in Finnsheep, Texel and F1. Gene names that were not part of Rambouillet reference transcriptome are marked with an asterisk (*) while gene IDs starting with LOC are “uncharacterized ncRNA”.

Similarly, *CA5A, FGF23* and *LOC114109527* (*NCLP*) were exclusively upregulated in F1 and *STX7* was downregulated. Among these, *CA5A* appeared to be upregulated in F1 crosses compared to both Finnsheep and Texel from both phases (i.e., in the CL in this study and in the ovary in our earlier study). *CA5A* is a member of the carbonic anhydrase family of zinc-containing metalloenzymes, whose primary function is to catalyze the reversible conversion of carbon dioxide to bicarbonate. The mitochondrial enzyme *CA5A* plays an important role in supplying bicarbonate (HCO_3-_) to numerous other mitochondrial enzymes. In a previous study, we observed downregulation of *CA5A* in the ovaries of Texel compared to F1 (Pokharel et al., 2018). More recently, *CA5A* was also shown to be expressed in the ovaries of the Pelibuey breed of sheep; the gene was upregulated in a subset of ewes that gave birth to two lambs compared to uniparous animals (Hernández-Montiel et al., 2019). However, there are no reports regarding the expression and function of *CA5A* in the CL. Based on the results from our current and earlier reports (Pokharel et al., 2018; Hernández-Montiel et al., 2019), *CA5A* appears to have an important function, at least until the preimplantation stage of reproduction. The level of expression in F1 crosses in the CL followed the same pattern as that in the ovary, which led us to conclude that *CA5A* is heritable and potentially an imprinted gene. Further experiments are needed to determine whether the gene is associated with high prolificacy.

### 3.7. Influence of flushing in gene expression

Four genes were significantly differentially expressed between Finnsheep and Texel of which three were upregulated in Texel. The only gene upregulated in Finnsheep was cytochrome c oxidase subunit 7c, mitochondrial (*COX7C*). The three genes significantly upregulated in Texel were *PAPPA2, LOC114116073* (ncRNA, uncharacterized) and *LOC114118704* (Golgi apparatus membrane protein *TVP23* homolog B pseudogene).

Out of 12 genes differentially expressed between Finnsheep and F1, five were upregulated in Finnsheep while seven were upregulated in F1. *TVP23* pseudogene was always downregulated in Finnsheep as a result of flushing diet. None of the six significantly differentially expressed genes were upregulated in Finnsheep suggesting null influence of flushing diet in this breed. Similarly, *HOXD10* (see XU 2014, important for regulation of embryonic implantation), *LOC105611004* (ncRNA, uncharacterized) and *LOC114116073* were upregulated in F1 compared to Finnsheep.

### 3.8. Limitations and thoughts for future studies

We acknowledge certain limitations of this study. With sequencing costs becoming increasingly inexpensive, increasing the sample size of each breed group would certainly add statistical power. Given that time-series experiments are not feasible with the same animal, sampling could be performed with a larger group of animals at different stages of pregnancy to obtain an overview of gene expression changes. One ovary from each ewe was removed earlier (Pokharel et al., 2018) and all the CLs for this study was collected from the remaining ovary. Therefore, there might be some impact due to possible negative feedback effects on overall gene expression. It should be noted that overall gene expression and, more specifically, differential expression between breeds is inherently a stochastic process; thus, there is always some level of bias caused by individual variation (Hansen et al., 2011). The results from breeding experiments have shown that productivity traits such as litter sizes may not carry on to F2 crosses (F1 x F1) and/or backcrosses. Therefore, future experiments that involve F2 crosses and backcrosses would provide more valuable findings related to prolificacy. Moreover, by doing a reciprocal cross experiment, we might be able to get insight into POE and measure the potential contribution of the Texel and Finnsheep in each cross. In addition, replicating such experiments in different environments would be relevant for breeding strategies to mitigate the effects of climate change. To minimize or alleviate noise from tissue heterogeneity, single-cell experiments may prove beneficial in future studies.

## 4. Conclusions

Our phenotypic records indicated lower embryo survival rate in Finnsheep compared to Texel. The relative scarcity of transcriptomic information about the CL means that its functional importance is underrated. We identified several key transcripts, including coding genes (producing mRNA) and noncoding genes, that are essential during early pregnancy. Functional analysis primarily based on literature searches and earlier studies revealed the significant roles of the most highly expressed genes in pregnancy recognition, implantation and placentation. F1 crosses were more closely related to Finnsheep than to Texel, as indicated by phenotypic and gene expression results that need to be validated with additional experiments (with F2 crosses and backcrosses). Several genes with potential importance during early pregnancy (including *SIGLEC14, SIGLEC8, MRP4*, and *CA5A*) were reported in the CL for the first time in any species. The results from this study show the importance of the immune system during early pregnancy and may even have greater significance to high prolificacy as revealed by significant upregulation of immune-related genes in Finnsheep. We also highlight the need for improved annotation of the sheep genome and emphasize that our data will certainly contribute to such improvement. Taken together, our data provide new information to aid in understanding the complex reproductive events during the preimplantation period in sheep and may also have implications for other ruminants (such as goats and cattle) and mammals, including humans.

## Author Contributions

Conceptualization, J.K., and M.H.L; methodology, K.P., and M.W.; software, K.P.; formal analysis, K.P.; investigation, K.P.; resources, J.K., M.H., and J.P.; data curation, K.P.; writing—original draft preparation, K.P.; writing—review and editing, K.P., J.P., and J.K; visualization, K.P.; supervision, J.K., and M.H.L.; project administration, J.K., and M.H.; funding acquisition, J.K., and M.H.L. All authors have read and agreed to the published version of the manuscript.

## Acknowledgments

We are grateful to Anu Tuomola for providing the experimental facilities on her farm in Urjala and to Kati Kaisajoki from Lallin Lammas Ltd. for providing the slaughtering facilities. We thank Johanna Rautiainen, Arja Seppälä, Annukka Numminen, Kalle Saastamoinen, Ilma Tapio, Tuula Marjatta Hamama, Anneli Virta and Magnus Andersson for their valuable assistance during this study. The authors wish to acknowledge the CSC – IT Center for Science, Finland, for computational resources. This study was supported by the Finnish Functional Genomics Centre of the University of Turku, Åbo Akademi and Biocenter Finland.

